# Radiance-Invariance Evaluation of a Compact LED-Based Source for Imager Radiance Transfer

**DOI:** 10.64898/2026.09.27.754831

**Authors:** Alberto J. Ruiz, Edwin A. Robledo, Eammon A. Littler

## Abstract

**Significance:** Fluorescence imaging remains largely qualitative and device-specific, limiting reproducibility and intersystem comparisons. SI-traceable radiometric imager characterization requires calibrated sources suitable for transferring known radiance to imaging systems.

**Aim:** Establish and evaluate a compact, calibrated solid-state radiometric emitter target (RET) for SI-traceable radiance transfer, and assess methods for evaluating radiance invariance, imager responsivity, and extension of radiance transfer to digitally defined regions of interest (ROIs).

**Approach:** The radiance invariance of an LED-based RET was evaluated by measuring its radiance across varying source-to-aperture distances and collection apertures. A previously described radiance-transfer formalism based on calibrated source radiance and collection geometry was implemented to determine imager responsivity (*R*_*λ*_), which was evaluated across varying working distances and lens apertures. Grubbs’ and MAD outlier tests and one-way ANOVA were used to assess radiance and responsivity invariance. Finally, responsivity obtained using digitally defined ROIs was compared with matched physical apertures to assess ROI-based radiance transfer.

**Results:** The measured RET radiance remained invariant across the tested source-to-aperture distances and collection apertures, with no significant differences observed across distance (p = 0.88) or aperture (p = 0.15). Imager responsivity remained stable across working distances for both tested lens configurations (p = 0.48 and p > 0.99). Responsivity was also invariant across lens apertures, except for deviations at f/11 that were consistent with practical aperture tolerances. Digitally defined ROIs reproduced responsivity values obtained using matched physical apertures, demonstrating that the radiance-transfer approach can be extended to image-defined regions.

**Conclusions:** The presented methods provide an approach for evaluating compact LED-based sources for invariant-radiance emission and suitability for SI-traceable radiance transfer. The characterized RET supported stable imager responsivity across changes in imaging geometry and extension to digitally defined ROIs. These methods can be adapted to other LED-based sources, with further angular characterization and uncertainty analysis needed to establish broader applicability.

## 1 Introduction

Fluorescence-guided imaging (FGI) and fluorescence-guided surgery (FGS) are increasingly used to provide molecular and physiological contrast during preclinical and clinical imaging; however, fluorescence measurements remain largely device-specific and qualitative.^1,2^ Imaging systems typically report device-native digital numbers rather than SI-traceable radiometric quantities, where the measured signal is influenced by collection geometry, optical throughput, detector response, acquisition settings, and image processing.^2,3^ Consequently, different imaging systems can report substantially different signals from the same fluorescent field, limiting quantitative comparison across devices, sites, and studies.^2–5^ Radiometric calibration provides a route to separate the measured optical signal from the device-specific response by relating image values to physical quantities such as fluorescence radiance (W·m^−2^·sr^−1^).^6–11^

Recent work by Litorja established an approach for transferring SI-traceable fluorescence radiance to an imaging system using calibrated radiance, known collection geometry, and conservation of étendue.^7^ In this approach, the radiant flux collected from a source of known radiance is related to the flux at the image plane, enabling determination of the imager responsivity and conversion of device-specific units to SI-traceable radiometric units (W·m^−2^·sr^−1^). An important practical requirement for this SI-unit transfer is a calibrated source that provides invariant emitted radiance over the collection geometries used for imaging.^12^ Spatial non-uniformities of a finite source, changes in source-to-aperture distance, or collection solid angle can result in geometry-dependent differences in the measured radiant flux and inferred radiance transfer. ^7,8^

Radiance standards are conventionally realized using calibrated lamps and integrating-sphere sources, which provide uniform diffuse output and invariant-radiance emission; these integrated-sphere sources are well established for radiometric calibration.^8–11^ However, the complexity and costs associated with these integrating-sphere sources that require the sphere, illumination source, and control/driving hardware have motivate the development of more compact and cost-effective form-factors.^13,14^ Advances in light-emitting diode (LED) technology make solid-state sources particularly suitable as the primary component of these calibrated emitters. This is further motivated by the fact that LEDs can provide relatively narrow spectral emission that can be selected to align with specific imaging bands, including to mimic the emission of fluorescence molecular agents; additionally, their well-characterized electrical-to-optical response enables reproducible electronic adjustment of radiant output. Solid-state LED sources have consequently been investigated as radiometric and low-radiant-flux standards, including LED sources developed for quantitative luminescence imaging.^13,15–19^ These characteristics provide a potential route toward compact, wavelength-specific radiance sources that can be readily adapted across fluorescence imaging bands without requiring a broadband source and subsequent spectral selection.

Despite these advances, the use of compact solid-state sources for radiance transfer remains limited. Methods are needed to establish whether such sources provide the radiance invariance^12^ required over relevant imaging geometries and whether they can reproduce the radiance-transfer behavior previously established using conventional reference sources.

Here, we characterized a calibrated solid-state radiometric emitter target (RET) and evaluated its radiance invariance across source-to-aperture distances and collection apertures. Subsequently, we implemented Litorja’s radiance-transfer framework^7^ to determine imager responsivity and evaluated its invariance across working distances and lens apertures. Finally, we compared matched physical and digitally defined apertures to assess the extension of radiance transfer to image-based regions of interest. Together, these experiments provide methods for evaluating solid-state sources for radiance transfer and assess their utility for SI-traceable radiometric characterization of imaging systems. To our knowledge, this study provides the first independent experimental evaluation of Litorja’s radiance-transfer formalism and extends this approach through direct evaluation of radiance invariance for a compact LED-based source and comparison of matched physical and digitally defined apertures for ROI-based radiance transfer.

## 2 Methods

### 2.1 Calibrated Solid-State Emitter: Radiometric emitter target

The evaluated calibrated solid-state emitter (radiometric emitter target, RET) is shown in **Fig 1** (QUEL Imaging, RRL-825-WR01-QUEL01) alongside the measured radiance vs. voltage curve, spectrum, and intensity profile characterizations. The RET consists of an LED, diffuser stack, aluminum base, and 3D printed housing components [**Fig 1(a)**]. The RET is voltage controlled to produce a deterministic radiance output; the model of RET used can provide an output radiance range of ~ 0.1–100 µW·cm^−2^·sr^−1^ [**Fig 1(b)**] with a measured spectral peak output centered at 827 nm with a full-width half-maximum of 35 nm. This RET output spectrum overlaps that of indocyanine green (ICG) in plasma [**Fig 1(c)**]. Although we used an 820-830 nm NIR band for ICG imaging applications, the presented calibration and evaluation methodologies are wavelength-agnostic. The RET exhibits a pseudo-flat emission profile [**Fig 1(c,d)**]. The RET calibrated radiance values were determined using **Equation (1)** from measurements obtained with a NIST-traceable photodiode (Thorlabs Inc., S120VC) positioned behind a 5 mm pinhole aperture (Thorlabs Inc., P5000K) positioned at 100 mm distance in a custom 3D-printed enclosure. This radiance characterization was performed using a 1.1-1.34 V driving-voltage range, which corresponded to a measured radiance emission range of 0.04 – 237 µW·cm^−2^·sr^−1^ [**Fig 1(b)**]. The radiance of the RET was set to ~50 µW·cm^−2^·sr^−1^ for the subsequent evaluations to provide sufficient measured signal for the tested distance and aperture ranges as described in the subsequent sections.

**Fig. 1:**
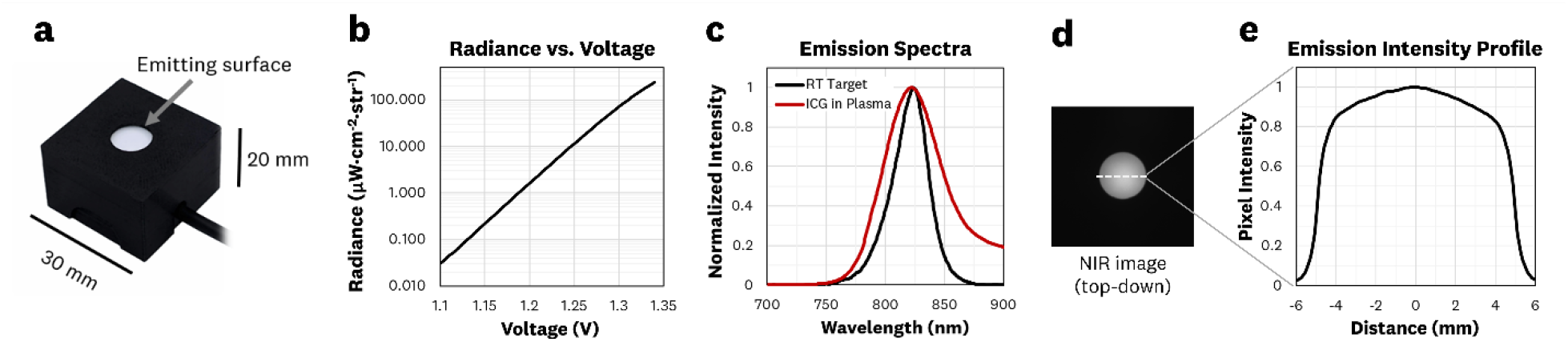
The RET alongside radiance, spectrum and intensity profile characterization: **(a)** The RET has a ~10 mm emitting output aperture with **(b)** voltage control to produce a radiance output of ~0.1-100 µW·cm^2^·sr. **(c)** The spectral output of the utilized RET falls within the spectra of ICG in plasma, indicating adequacy for use with most 800 nm channel fluorescence systems. **(d)** The RET has a pseudo-flat emission profile providing a diffuse output for imaging applications.

### 2.2 Evaluating the RET radiance invariance with distance and collection solid angle

To enable the conversion of imager response to SI units, the RET must exhibit radiance-invariant behavior.^8^ In brief, this means that radiance must remain invariant with distance and solid angle, enabling direct mapping of an SI-traceable radiance (W·m^−2^·sr^−1^) to the imager response.^7,8^ The radiance emitted by a source (*L*_*src*_) can be determined by the detected power from a calibrated photodiode power sensor (Φ_*ps*_) for a given emitter source area (*A*_*src*_) and associated solid angle (Ω_*src*_). As shown in **Fig 2(a)**, Ω_*src*_ is defined by the aperture stop (*A*_*apt*_) and the source-to-aperture distance (*d*). Mathematically the radiance is defined as

**Fig. 2:**
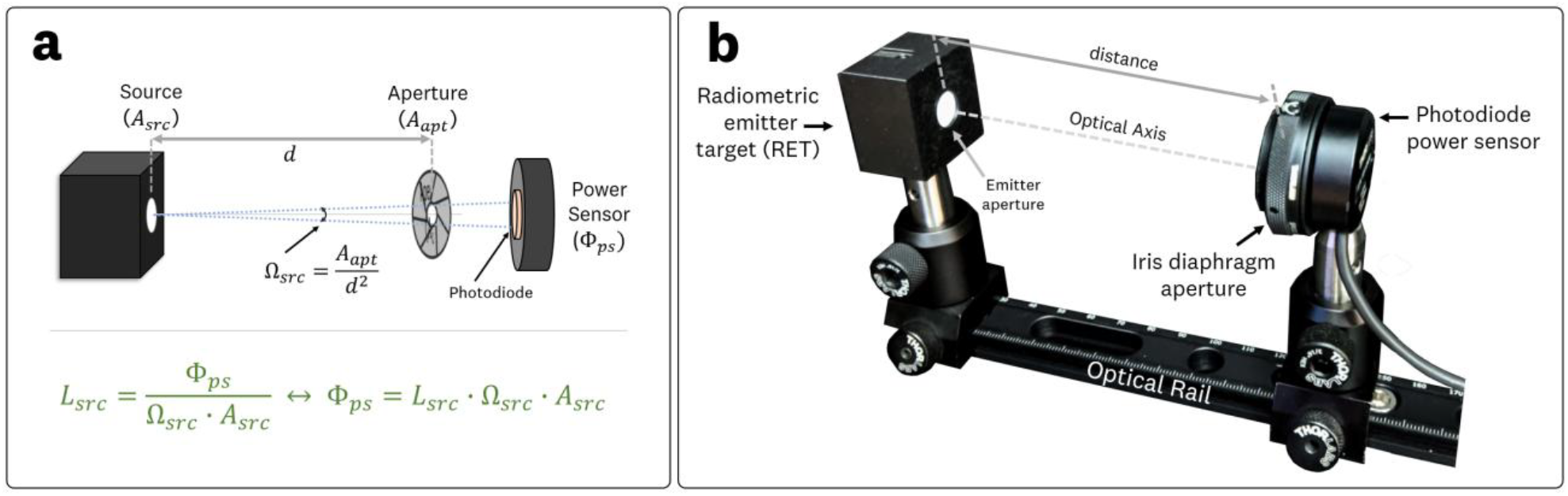
Optical set-up for assessing the RET’s radiance-invariant behavior: **(a)** Geometric optics schematic for radiance *L*_*src*_ measurement, illustrating the light transmission from source to detector alongside relevant quantities and **(b)** experimental set-up for testing the radiance-invariant behavior of the RET.

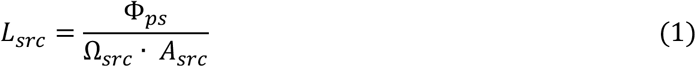

Where 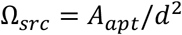. Conversely, if the radiance is known, the detected radiant flux (i.e. power) is given by

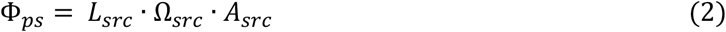

These mathematical relationships indicate that a calibrated photodiode can be used to measure the radiance output of a source, by additionally measuring *d, A*_*src*_, and *A*_*apt*_.

**Fig 2(b)** shows the experimental setup used for assessing the RET’s radiance invariance, which utilized a calibrated photodiode (Thorlabs Inc., S120VC) and a variable iris diaphragm aperture (Thorlabs Inc., SM1D12C) mounted on an adjustable optical rail. The RET and photodiode were aligned along the optical axis at a *d* = 400 mm to achieve consistent optical power measurements within the tested 25 - 400 mm range. To test the radiance invariance with distance, the aperture was fixed at 5.0 mm (*A*_*apt*_ =19.6 mm^2^) and the detected power recorded at *d* = 25, 50, 75, 100, 150, 200, 250, 300, 350, and 400 mm. To test the solid-angle invariance, the distance was fixed at *d* = 100 mm, while the aperture diameter varied at 2, 3, 4, 5, 6, 7, and 8 mm utilizing the markings of the variable iris diaphragm aperture. Each measurement series was acquired in triplicate. Furthermore, the true aperture diameter at these diaphragm markings were measured in triplicates using a calibrated caliper (Mitutoyo, 500-196-30), such that the aperture diameters used in the radiance calculation were of 1.86, 2.83, 3.90, 4.93, 5.98, 6.99, and 8.05 mm.

#### 2.2.1 Statistical tests for outliers and group differences

Under the radiance-invariant behavior, radiance should not vary with distance or aperture. To evaluate this, statistical analyses were performed to identify outliers and group mean differences across each condition. Outlier detection was performed using Grubbs’ test (one-sided, *α*=0.05) and a Median Absolute Deviation (MAD) rule of thumb (using *k*=3 as the cutoff). For Grubbs’ test, the test statistic was compared to the critical value *G*_*critical*_, for *n* datapoints at *α*=0.05. A data point was flagged as an outlier if *G > G*_*critical*_. Separately, the MAD approach computed each point’s absolute deviation from the median. Any point whose deviation exceeded *k* times the robust deviation estimate was labeled as an outlier. To evaluate whether mean radiances differed significantly across distance or aperture settings, we performed a one-way ANOVA at *α* = 0.05. Each distance or aperture condition formed one “group,” with repeated measurements (triplicates) providing within-group variance. The resulting *F*-ratio and associated *p*-value were used to test the null hypothesis that all group means were equal.

### 2.3 Conversion of radiance into camera response

This section recreates the radiometric conversion formalism of Litorja^7^ to validate that the RET can serve as a calibrated source for SI-traceable radiance transfer under standard imaging geometries. Using the RET’s calibrated radiance output for conversion of the camera response involves imaging the RET emission aperture while accounting for the lens aperture size and the source-to-aperture distance. The relevant parameters and simplified 2D geometric schematics involved in the imager response conversion are shown in **Fig 3(a,b)**. The following derivations and equations are adapted from Litorja.^7^ Two optical geometries are considered: (1) imaging of the RET emission area onto the image sensor [**Fig 3(a)**] and (2) the aperture-limited light rays for the projection of a single source emission point onto the sensor [**Fig 3(b)**]. These two configurations define four solid angles determined by the source area (*A*_*src*_), image area (*A*_*img*_), aperture size (*A*_*apt*_), source-to-aperture distance (*d*) and aperture-to-image sensor distance (*d*_*sen*_). In the first geometry [**Fig 3a**] the two defined solid angles are 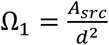 and 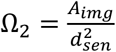 where Ω = Ω. For the second optical geometry [**Fig 3b**] the two defined solid angles are 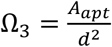 and 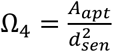. Note that the “source-to-aperture distance” *d* is measured from the emitting surface of the RET to the lens’s entrance pupil – the effective aperture position that defines the collection solid angle (Ω_3_) – rather than to the mechanical diaphragm location.

**Fig. 3:**
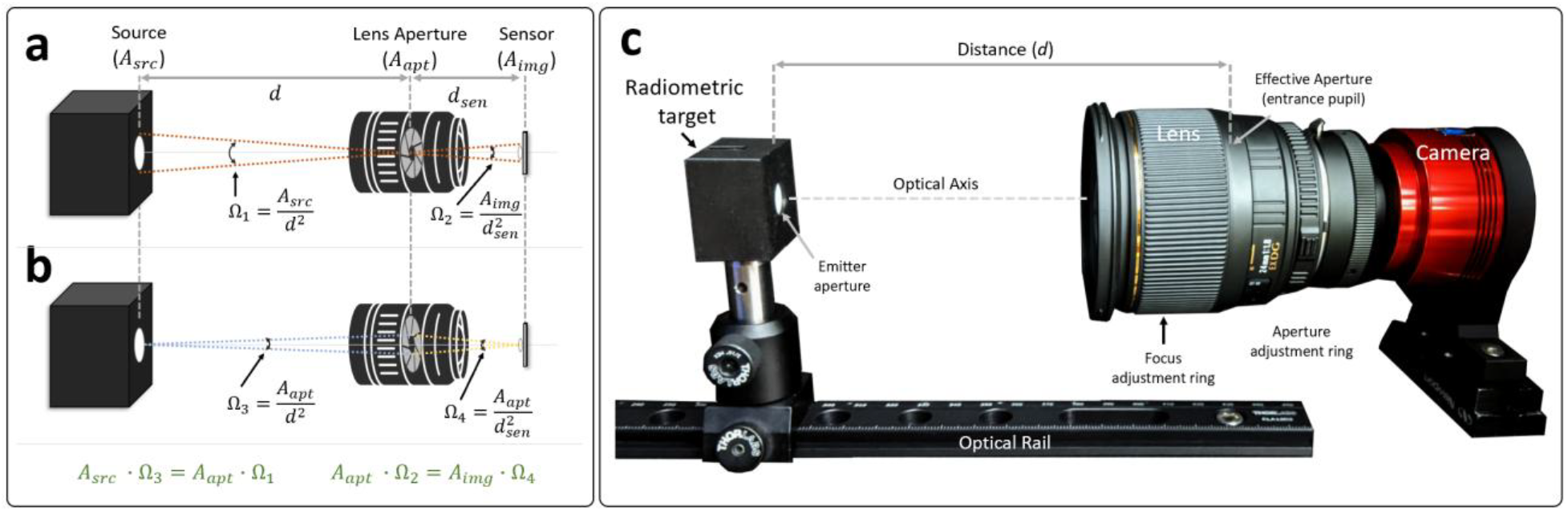
Optical set-up for assessing the radiance conversion of the camera response: **(a)** Geometry associated with imaging the RET source area onto the image sensor. **(b)** Geometry associated with the aperture-limited light rays for the projection of a single source emission point onto the sensor. **(c)** Experimental setup used to test the imager response conversion for varying distances and apertures.

As described in Litorja,^7^ the product of the area and solid angle *A* · Ω, often referred to as the *étendue*,^20^ remains constant in an optical system. This principle enables calculation of the radiant flux (Φ) passing from the RET through the lens aperture and projected onto the imaging sensor. The radiant flux, which is reduced by the lens transmittance, is projected onto the imaging sensor, such that:

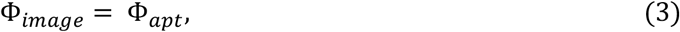

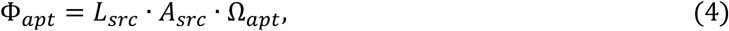

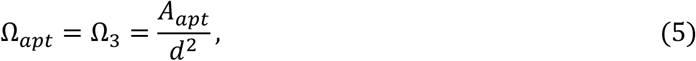

So that

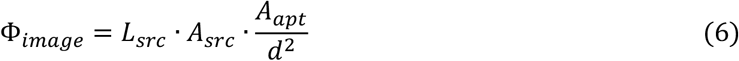

Hence, the measured radiant flux at the image plane *Φ*_*image*_ can be determined from the RET radiance (*L*_*src*_), RET source area (*A*_*src*_), lens aperture area (*A*_*apt*_), and source-to-aperture distance (*d*). From an imaging perspective, photons emitted by the RET are converted to electrons within each pixel, then digitized to a corresponding digital number *N* at each pixel (bound by the camera bit depth). The sum of the digitized outputs (*N*_*i*_) for all pixels (*i*) corresponding to the RET emitting area is noted as *S*, such that

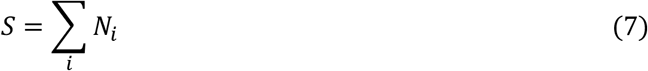

Lastly, the imager responsivity in the specified spectral band, *R*_*λ*_, is the ratio of total counts to the image-plane radiant flux:

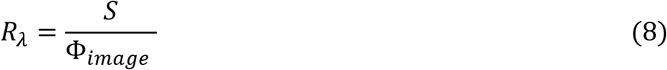

with units of counts/W. Hence, if the source exhibits radiance-invariance and the principle of *étendue* holds, *R*_*λ*_ should remain invariant with both distance and aperture size – thereby enabling direct conversion of SI-traceable radiance into the camera response.

#### 2.3.1 Imager Responsivity vs. Distance

**Fig 3(c)** shows the experimental setup used to test imager responsivity *R*_*λ*_ with varying source-to-aperture distances (*d*). A 12-bit back-illuminated CMOS camera (Suzhou ZWO Co, ASI462MM) and lenses were used to image the RET emission surface with the RET optically aligned on an adjustable optical rail. The RAW image data (12-bit native digitization) were exported in 16-bit TIFF format and used for subsequent analysis. A Navitar 16mm lens (Navitar, Machine Vision Lens 16mm f1.4) and Sigma 24 mm lens (SIGMA Corporation, #432306) were used to test the imaged radiance invariance with distance by using a fixed aperture area (*A*_*apt*_) and varying the source-to-aperture distance (*d*). The Navitar images were acquired with a fixed f-number aperture of 4.0, gain of 80, exposure time of 10 ms, and *d* = 100, 150, 200, 300, 350, and 400 mm; the Sigma lens images were acquired with a fixed f-number aperture of 5.6, gain of 80, exposure time of 10 ms and *d* = 100, 150, 200, 300, and 350 mm. The exposure times were selected to ensure similar signal-to-noise performance and keep the imaging sensor within its linearity range (i.e., ~80% of total well capacity). Images were acquired with a RET radiance output of 54 µW·cm^−2^·sr^−1^. Aperture size was determined by each lenses’ f-number *n* setting, defined as *n* = *f*/*D*, where *f* is the focal length of the lens and *D* is the aperture diameter. The corresponding aperture area *A*_*apt*_ can be calculated from:

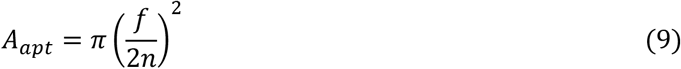

Therefore, the calculated aperture area *A*_*apt*_ for the Sigma and Navitar lens acquisitions were 14.4 mm^2^ and 12.6 mm^2^, respectively. Each lens was refocused at every tested *d* to ensure the RET emission surface remained in focus, thereby preserving image quality and measurement consistency across distances.

#### 2.3.2 Imager Responsivity vs. Aperture Size

To examine the invariance of the imager responsivity *R*_*λ*_ with respect to aperture settings, the aperture size (*A*_*apt*_) was varied at a fixed source-to-aperture distance (*d*). The Sigma 24mm lens and a Nikon 50mm lens (Nikon, #1433) were used. Both image sets were acquired with a RET radiance output of 54 µW·cm^−2^·sr^−1^ and f-number apertures settings of 2.8, 4, 5.6, 8, and 11. A fixed distance *d* of 300 mm and 400 mm were used for the Sigma and Nikon lenses, respectively. Imaging parameters were set to an exposure time of 9 ms and gain of 80, ensuring that the largest aperture setting (f/2.8) produced ~80% sensor saturation to preserve signal linearity. Each corresponding aperture area was calculated using **Equation (9)**.

#### 2.3.3 Image and Responsivity Analysis

All images (12-bit native digitization) were acquired as uncompressed 16-bit TIFFs and analyzed in ImageJ to extract the total pixel counts (*S*) over the 10 mm RET emission surface. Imaged radiant flux values Φ_*image*_ were calculated using **Equation (6)**. The resulting *R*_*λ*_ was calculated using **Equation (8)**. To evaluate whether the mean imager responsivity differed significantly across distance or aperture settings, we performed a one-way ANOVA at *α* = 0.05. Each distance or aperture condition formed one “group,” with repeated measurements (triplicates) providing within-group variance. The resulting *F*-ratio and associated *p*-value were used to test the null hypothesis that all group means were equal.

### 2.4 Physical vs. Digital apertures for ROÍ radiance transfer

To extend the concept of imager responsivity (*R*_*λ*_) to digitally defined region-of-interest (ROI) measurements, we evaluated the *R*_*λ*_ for both physical apertures and “digital apertures.” For this, the Sigma and Nikon lenses were used to image the RET under three physical aperture conditions: (1) No aperture (RET emission surface of 10 mm ∅), a 5 mm physical aperture (Thorlabs, P5000K), and 2 mm physical aperture (Thorlabs, P2000K). All images were acquired using a RET radiance of 54 µW·cm^−2^·sr^−1^, camera gain of 80, an integration time of 30 ms, and an f-number of 5.6, with the Sigma lens positioned at 300 mm and the Nikon lens at 400 mm. The ZWO camera was used for imaging. By applying digital masks (ROIs) to these images in post-processing, the variability of *R*_*λ*_ with both physical aperture size and smaller digitally defined regions was assessed. All images (12-bit native digitization) were acquired as uncompressed 16-bit TIFFs and analyzed in ImageJ.

## 3 Results

### 3.1 Radiance-invariant emission of the radiometric target

The results of evaluating the RET’s radiance-invariant emission are shown in **Fig 4**, which compares measured radiance under two conditions: (a) varying source-to-aperture distance and (b) varying aperture size. Statistical analyses included Grubbs’ and Median Absolute Deviation (MAD) outlier tests, as well as one-way ANOVA across each set of mean radiance values. The methods of these measurements and analysis are covered in Section 2.2.

**Fig. 4.**
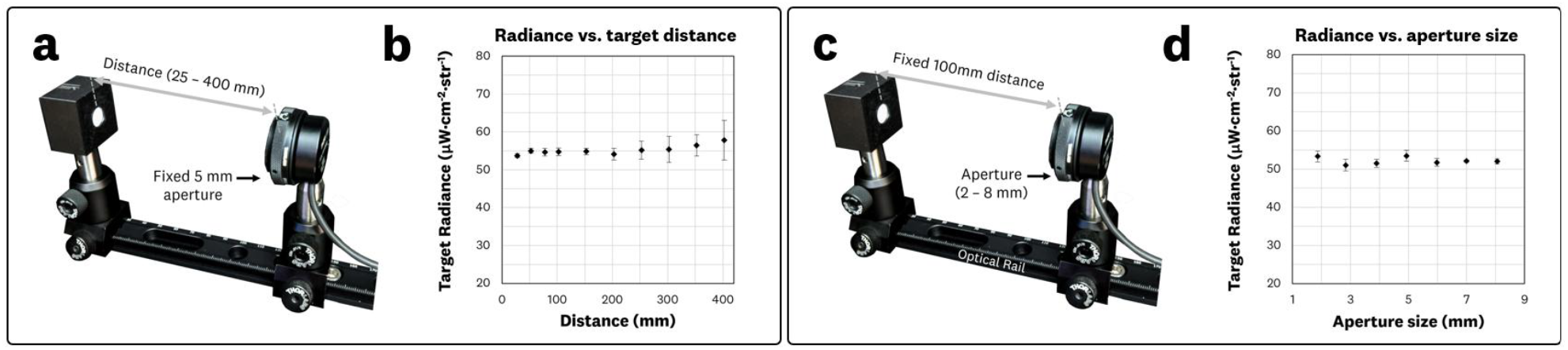
Results of the radiance-invariance evaluation of the RET: **(a)** varying distance experimental set-up and (b) plot of the measured radiance vs. distance data. **(c)** Varying aperture experimental set-up and **(d)** plot of the measured radiance vs. aperture size data.

In the distance-varying experiment [**Fig 4(a,b)**], no outlier was flagged by Grubbs’ test (*n=*10, *G*_*max*_ = 2.28 < *G*_*critical*_ = 2.82). However, the largest distance (400 mm) was identified as an outlier by the MAD criterion (400 mm radiance = 57.7 > cut-off value = 56.7 µW·cm^−2^·sr^−1^), likely due to the signal approaching the detector’s lower detection limit at this distance. Additionally, a trend of increasing standard deviation was observed as distance increased, corresponding to a decline in signal-to-noise ratio; for instance, the 400 mm measurement had approximately 200× less signal than the 25 mm measurement. Despite this discrepancy, a one-way ANOVA across all tested distances showed no significant difference (p ≈ 0.88), indicating radiance remained effectively constant with distance.

For the aperture-varying experiment [**Fig 4(c.d)**], neither Grubbs’ test (*n=*7, *G*_*max*_ = 1.43 < *G*_*critical*_ = 2.02) nor MAD criteria (max deviation = 1.47 < MAD cut-off = 2.38 µW·cm^−2^·sr^−1^) flagged any outlier. A one-way ANOVA also did not reveal any statistically significant difference (*p* = 0.15) among aperture settings. These findings confirm that the measured radiance remained effectively constant with the solid angle subtended by the aperture.

Overall, the combined outlier and ANOVA results demonstrate that the RET emission remains consistent over the tested distances and apertures. This radiance-invariant behavior provides the basis for converting imager response into SI-traceable units.

### 3.2 Conversion of radiance to camera response

Having established the radiance-invariant behavior of the RET (Section 3.1), we next evaluated the conversion of radiance into imager response according to the methods in Sections 2.3, 2.3.1, and 2.3.2. Specifically, we computed the imager responsivity, *R*_*λ*_, using **Equations (6)-(8)** for different source-to-aperture distances (*d*) and for varying aperture sizes (*A*_*apt*_). **Fig 5** and **Fig 6** summarize these results.

**Fig. 5.**
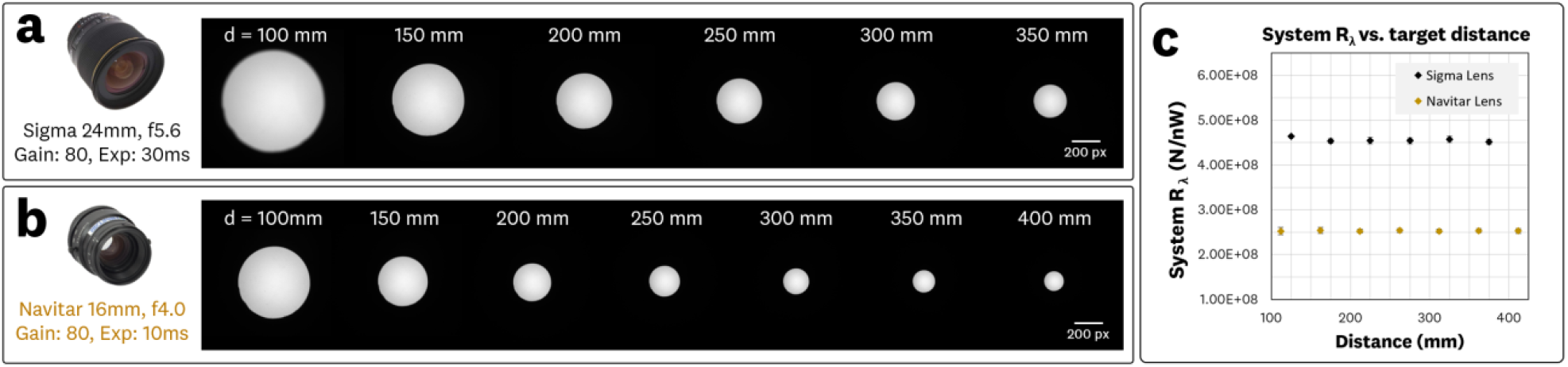
Results of the imager responsivity *R*_*λ*_ vs. source-to-aperture distance *d* measurements: **(a)** Sigma lens varying distance acquired images, **(b)** Navitar lens varying distance acquired images, and (c) the plot of the resulting system responsivity *R*_*λ*_ vs. source-to-aperture distance *d* measurements. All images share the same look-up-table scaling to provide cross-image comparison.

**Fig. 6.**
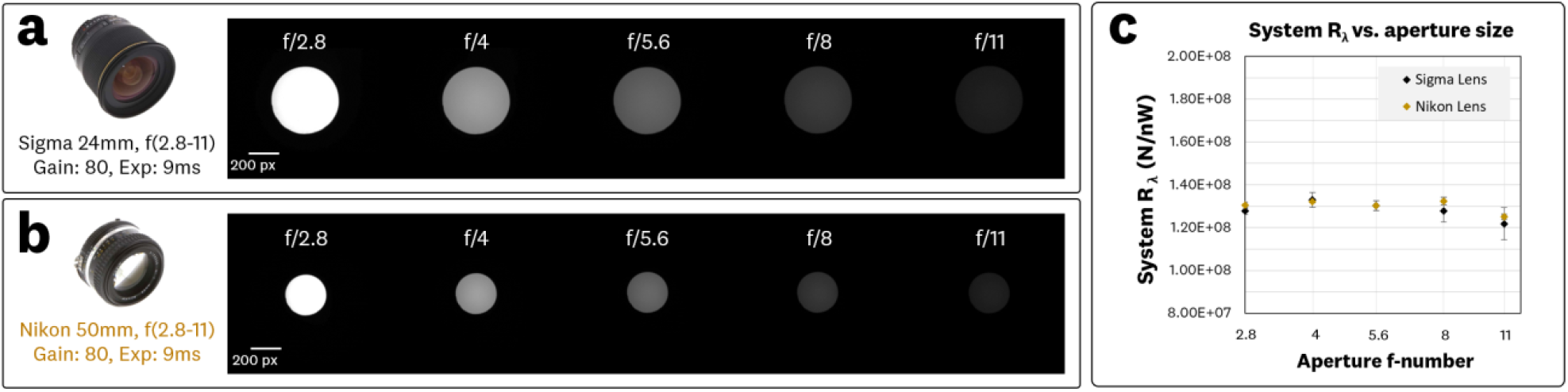
Results of the imager responsivity *R*_*λ*_ vs. aperture size comparison: **(a)** Sigma lens varying aperture acquired images, **(b)** Nikon lens varying aperture acquired images, and **(c)** the plot of the resulting system responsivity *R*_*λ*_ vs. aperture f-number plots measurements. All images share the same look-up-table scaling to provide cross-image comparison

#### 3.3.1 Imager responsivity vs. distance

**Fig 5** shows the acquired images for the Sigma 24 mm lens [**Fig 5(a)**] and Navitar 16 mm lens [**Fig 5(b)**] alongside the plotted *R*_*λ*_ vs *d* data [**Fig 5(c)**]. All images were displayed using a common look-up-table scaling (0–16-bit range mapped to black and white, respectively) to enable direct cross-image comparison. Both lenses were used to image the RET at distances *d* of 100–400 mm with fixed apertures. The measured *R*_*λ*_ remained consistent across all tested distances, with average values of 4.47 counts/W [**Fig. 5(a)**] and 2.48 counts/W [**Fig. 5(b)**] for the Sigma and Navitar lens acquisitions, respectively. A one-way ANOVA indicated no statistically significant difference among these mean values (p = 0.48 for the Sigma lens; *p* > 0.99 for the Navitar lens). These findings confirm that, under the tested conditions, the imager response scales appropriately with the RET radiance regardless of distance, in line with the radiance invariance and *étendue* assumptions.

The difference in the absolute *R*_*λ*_ between the two lenses reflects variations in imaging parameters (i.e., aperture setting, exposure time) as well as differences in lens transmission and optical design. Nonetheless, each system’s *R*_*λ*_ remained stable over the examined distance range, supporting the expected invariance of responsivity with distance for radiance-invariant source.

#### 3.2.2 Imager responsivity vs. aperture size

**Fig 6** shows the acquired images for the Sigma [**Fig 6(a)**] and Nikon [**Fig 6(b)**] lenses alongside the plotted *R*_*λ*_ vs aperture size data [**Fig 6(c)**]. All images share the same look-up-table scaling to provide cross-image comparison; the scaling used was normalized to the imaging set maxima. Identical camera settings were utilized with fixed distances and varied apertures as described in Section 2.3.2.

For the Sigma Lens dataset [**Fig 6(a)**], a one-way ANOVA provided *p* = 0.10 when all five apertures measurements were included, indicating no statistically significant difference in responsivity. Excluding the f/11 measurement (where small manufacturing tolerances can have a larger relative impact) increased *p* to 0.26. For the Nikon Lens dataset [**Fig 6(b)**], the ANOVA provided *p* = 0.001 when all f-numbers were included, but rose to 0.45 when the f/11 datapoint was removed. This improvement suggests that the largest f-number measurement can be sensitive to minor lens-aperture deviations (e.g., manufacturing tolerances of ±100 µm) that become more proportionally significant at smaller aperture diameters; these manufacturing deviations, along with the inability to measure the true lens aperture size, can alter the statistical outcome of the ANOVA test. Similar observations have been reported previously by Litorja.^7^

Overall, these aperture-variation results are consistent with the theoretical expectation that, under radiance invariant emission and *étendue* assumptions, *R*_*λ*_ should not depend on the lens aperture. Where minor deviations did arise (i.e., aperture settings of f/11), they appear attributable to practical manufacturing tolerances rather than a violation of the *étendue* principle.

### 3.3 Physical vs. Digital apertures for radiance transfer

Having confirmed the radiance-invariance behavior of the RET (Section 3.1) and validated the conversion of radiance to imager response (Section 3.2), we next examined whether digital region-of-interest (ROI) selection could reliably replace physical apertures when determining the imager responsivity *R*_*λ*_. This approach is relevant to workflows where ROI-based measurements are used to isolate a sub-area of the imaged field. Unless otherwise noted, all *R*_*λ*_ values in this section are expressed in units of x10^8^ counts/nW.

**Fig 7(a)** shows representative images of the RET acquired with no physical aperture (10 mm emission area), a 5 mm physical aperture, and equivalent digital ROI. **Fig 7(b)** and **Fig 7(c)** present the resulting *R*_*λ*_ values for the Sigma and Nikon lens imaging sets, respectively. For the Sigma lens set [**Fig 7(b)**], the measured *R*_*λ*_ for the physical vs. digital 5 mm aperture was (4.8 ± 0.1) vs. (4.8 ± 0.1), and for the 2 mm aperture (5.0 ± 0.3) vs. (5.0 ± 0.1). Similarly the Nikon lens set [**Fig 7(c)**] yielded (5.1 ± 0.1) vs. (5.1 ± 0.1) for the 5 mm aperture and (5.2 ± 0.2) vs. (5.3 ± 0.1) for the 2 mm aperture. Given the difference in mean responsivity values for the 2 mm Nikon data, a two-tailed Welch t-test was performed (independent samples; N = 1,800 pixels; assumed spatial correlation factor = 0.05), which yielded *p* = 0.32, indicating no statistically significant difference in responsivity. These results indicate that there was no statistical difference between physically apertured measurements and those obtained via digital masking, which signifies that adjacent illuminated regions did not contribute substantially to the ROI’s measured radiance.

**Fig 7.**
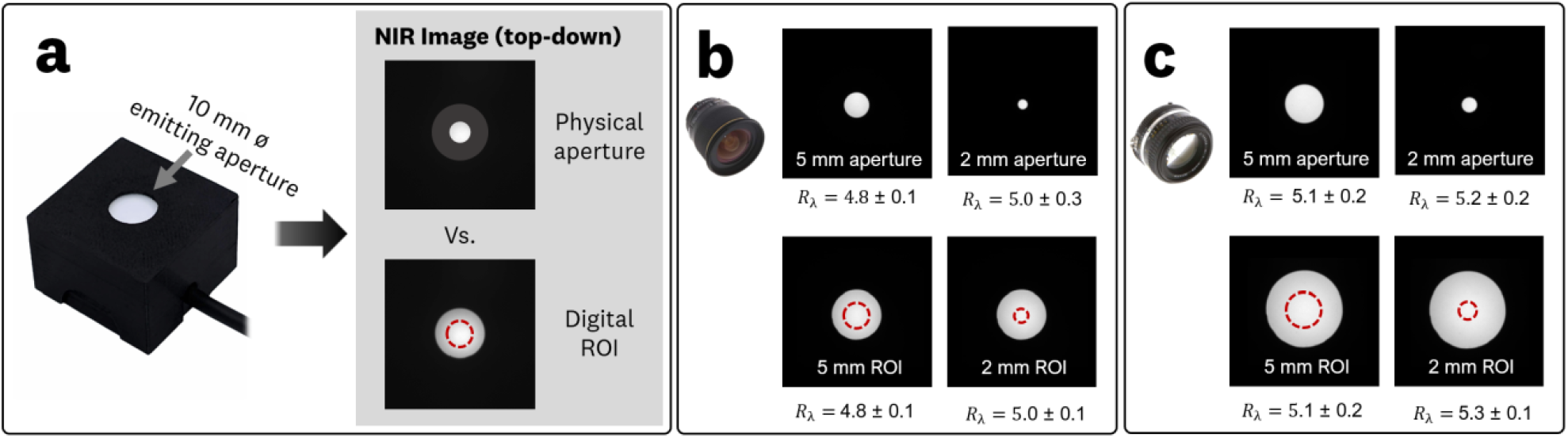
**(a)** Physical vs. digital ROI system responsivity experiment utilizing 5 mm and 2 mm apertures with respective images and calculated *R*_*λ*_ values for the imaging configurations utilizing the **(b)** Sigma 24 mm lens, and **(c)** Nikon 50 mm lens.

For the full 10 mm emission region, the measured *R*_*λ*_ values were 4.4 ± 0.3 (Sigma image) and 4.7 ± 0.4 (Nikon image). These values are slightly lower than those for the smaller apertures, reflecting the pseudo top-hat emission profile of the RET [**Fig 1(e)**]: the smaller central regions tend to have a more uniform, slightly higher radiance than the periphery. Although this difference does not alter the conclusion that physical and digital apertures provide equivalent results, it highlights the need to account for potential non-uniformities in the calibrated solid-state emitter when deriving *R*_*λ*_. This indicates that, in practice, responsivity values tied to the RET’s calibrated radiance *L*_*cal*_ should be derived from total digital counts *S* over the full 10 mm ROI to account for the pseudo-top-hat profile and minimize bias from peripheral roll-off [**Equations (7),(8)**].

## 4 Discussion

In this work, we evaluated the radiance invariance of a calibrated solid-state radiometric emitter target (RET) and independently implemented Litorja’s radiance-transfer framework^7^ for SI-traceable radiance transfer. The measured RET radiance remained invariant across the tested source-to-aperture distances and collection apertures, while imager responsivity (*R*_*λ*_) remained stable across varying working distances and lens apertures. Digital ROIs also reproduced responsivity values obtained using matched physical apertures, demonstrating that the radiance-transfer approach can be extended to digitally defined image regions. The subsections below further discuss the results.

### 4.1 Radiance-invariant emission of the radiometric target

The radiance-versus-distance and radiance-versus-aperture experiments (Section 3.1, **Fig 4**) confirmed that the RET exhibited radiance-invariant emission for this on-axis orthogonal imaging. Across both experimental series, measured radiance showed no statistically significant dependence on distance or aperture, as supported by ANOVA results. Minor discrepancies at the largest working distance (400 mm) were attributed to reduced signal-to-noise, which in turn led to marginal outlier identification by robust statistical tests, but these did not alter the overall conclusion. Importantly, radiance remained constant within experimental uncertainty, providing empirical confirmation of radiance-invariant emission. This invariance is essential for establishing SI-traceable radiance transfer.

### 4.2 Conversion of radiance to camera response

The imager responsivity experiments (Section 3.2, **Fig 5,6**) further confirmed the radiance-invariant emission of the RET and its suitability for imager response conversion and characterization. The results indicate that imager responsivity (*R*_*λ*_) was invariant, within experimental uncertainty, across the tested working distances and f-numbers, as expected from the radiance-invariant emission and *étendue* conservation described in Eqs. (3)–(8). This invariance validates the theoretical requirement for converting camera-specific responses into SI-traceable radiometric units. The *R*_*λ*_ departures observed at the smallest apertures (f/11) are most likely attributed to manufacturing tolerances of the aperture, which has been previously reported;^7^ for example, a 100 µm deviation for an f/11 aperture setting would be sufficient to account for the observed deviations. The differences in *R*_*λ*_ observed between lens/camera configurations are consistent with expected contributions from varying lens transmission, effective f-number, and acquisition parameters (exposure time, gain). Importantly, while a distance- and aperture-independent *R*_*λ*_ provides a local linear mapping from digital counts (*N*) to calibrated radiance (*L*_*cal*_) at a given operating point, it does not account for sensor or image-processing nonlinearities across the full dynamic range, motivating the development and evaluation of the RTC.

### 4.3 Physical vs. Digital apertures for radiance transfer

The physical-versus-digital aperture experiments (Section 3.3, **Fig 7**) showed that ROI masking produced imager response (*R*_*λ*_) values statistically indistinguishable from those obtained with matched physical apertures. This confirms that adjacent illuminated regions did not statistically contribute to the ROI signal, enabling digital ROIs to substitute for physical apertures in responsivity estimation. Slightly lower *R*_*λ*_ values observed for the full 10 mm emission area reflect the RET’s pseudo top-hat profile, underscoring the need to account for spatial non-uniformity when extending calibrations from small ROIs to the full emission surface. Practically, this means that responsivity measurements utilizing the RET should use the entire emission surface as the ROI to minimize the impact of spatial non-uniformity. Overall, these results demonstrate that digital ROI selection is a valid approach for radiometric calibration and directly supports the ROI-level radiance transfer. These results also inform the future exploration of pixel-level radiance transfer given the discrete ROIs provided by camera sensor pixels/photosites.

### 4.6 Limitations and Future Work

Limitations in this study center around the exclusive characterization of the RET under an on-axis geometry; while the distance- and aperture-invariance results support its use for the geometries evaluated here, its angular radiance distribution was not characterized. Off-axis measurements and angular emission characterization are therefore needed to establish performance across broader viewing geometries, including understanding if the radiance-invariance extends to the angular behavior of Lambertian sources.^8,18^ Furthermore, a formal uncertainty budget for RET radiance and transferred imager responsivity should incorporate contributions from radiance calibration, aperture and entrance-pupil geometry, detector response, and source non-uniformity. Extending characterization to additional spectral bands and imaging systems would further help establish the robustness and generalizability of the radiance-transfer approach.

The radiance-transfer formalism utilized here, consistent with previous approaches,^7,9^ characterizes imager response at discrete radiance operating points and therefore does not fully account for sensor nonlinearities or transformations introduced by sensor response non-linearities and image-processing pipelines. The limitations imposed by these established approaches is studied in subsequent work from our group, where extended the radiance-invariant RET approach through the introduction of the Radiance Transfer Curve (RTC), Radiance Imaging Transform (RIT), and Fluorescence Imaging Transform (FIT) to characterize response across the camera sensor dynamic range, address image-processing pipelines, and enable pixel-level radiometric and excitation-normalized fluorescence imaging.^21^

## 5 Conclusion

This study demonstrates an approach for evaluating the radiance invariance of a compact, calibrated LED-based source across source-to-aperture distances and collection apertures under defined imaging geometries. Importantly, these methods can be adapted to other radiance calibration sources to evaluate radiance invariance and establish their suitability for radiance-transfer applications. Using the characterized RET, we performed the first independent implementation of Litorja’s radiance-transfer framework^7^ and demonstrated invariant imager responsivity across changes in working distance and lens apertures. The agreement between matched physical and digital aperture measurements further demonstrated that the radiance-transfer approach can be extended to digitally defined ROIs for responsivity estimation, along the potential expansion to pixel-level radiance transfer. Together, these results provide a practical basis for evaluating compact calibrated sources and their use for SI-traceable radiometric characterization of imaging systems. Further angular characterization, formal uncertainty analysis, and evaluation across additional imaging systems and spectral bands will be important for establishing the broader applicability of these sources for SI-traceable radiometric characterization.

## Disclosures

A.J.R. is the co-founder and chief technology officer of QUEL Imaging. E.A.R. is employed full-time by QUEL Imaging. E.A.L. is employed full-time by QUEL Imaging. QUEL Imaging designs, manufactures, and supplies reference targets, phantoms, and tools to support the development lifecycle of fluorescence imaging systems and other optical technologies.

## Acknowledgments

This project was funded in part with federal funds from the National Institute of Biomedical Imaging and Bioengineering, National Institutes of Health, Department of Health and Human Services (Grant No. R43/44EB029804). The breast lumpectomy model utilized in this study was funded by the National Institutes of Health National Cancer Institute (NCI) contract 75N91021C00035.

The authors also gratefully acknowledge Toni Litorja for her valuable guidance and discussions related to the radiance transfer and radiometric calibration aspects of this work.

The authors also acknowledge the use of the ChatGPT-5 language model (OpenAI), Writefull (https://www.writefull.com), and the Microsoft Word Editor tools for editorial editing, grammar correction, and text refinement.

## Code and Data Availability

The data used in this study are available from the corresponding author upon reasonable request.

